# InterPep2: Global Peptide-Protein Docking with Structural Templates

**DOI:** 10.1101/813238

**Authors:** Isak Johansson-Åkhe, Claudio Mirabello, Björn Wallner

**Affiliations:** Linköpings Universitet; Linköping University

## Abstract

**Motivation:** Interactions between proteins and peptides or peptide-like intrinsically disordered regions are involved in many important biological processes, such as gene expression and cell life-cycle regulation. Experimentally determining the structure of such interactions is time-consuming, and because of the disordered nature of the ligand, the interactions are especially difficult to predict through software, requiring specialized solutions. Although several prediction-methods exist, most are limited in performance or availability.

**Results:** InterPep2 is a freely available method for predicting the structure of peptide-protein interactions. We have previously shown that structural templates can be used to accurately predict peptide-protein binding sites, and that using templates from regular protein-protein interactions will increase the number of sites found. Here, we show that the same principle can be extended to dock the peptide to the binding surface using InterPep2. A key component of InterPep2 is the ability to score plausible interaction templates using a RandomForest trained to predict the DockQ-score using sequence and structural features. InterPep2 is tested on a difficult dataset of 251 peptide-protein complexes, where it correctly positions 136 (54%) at the correct site compared to 114 (45%) for the second best method. Analyzing the confidence score InterPep2 recalls more true positives across all specificity levels compared to the second best method, for example at 10% False Positive Rate it correctly identifies 59% of the complexes compared to 44% for the second best method.

**Availability:** The program is available from: http://wallnerlab.org/InterPep

**Contact:** Björn Wallner

## 1 Introduction

Protein-protein interactions are vital in most biological processes, from metabolism to cell life-cycle (Midic *et al.*, 2009; Tu *et al.*, 2015). To understand these processes, it is important to know the structural details of the interactions. Structures of interacting proteins can be experimentally solved through a multitude of methods such as x-ray crystallography, NMR, and cryo-EM (Rhodes, 2010; Wüthrich, 1986; Topf *et al.*, 2008). However, because of the complexity, cost, and time it takes to perform experiments, computational methods have been developed to support and supplement. Methods such as HADDOCK, PIPER, and ZDOCK (Dominguez *et al.*, 2003; Kozakov *et al.*, 2006; Pierce *et al.*, 2014) make predictions directly from sequence or energy function information, while others such as InterPred, PRISM, and M-TASSER (Wallner and Mirabello, 2017; Baspinar *et al.*, 2014; Chen and Skolnick, 2008) use already solved structures as templates for prediction. Among these methods, template-based approaches have shown great success in the past.

A significant fraction (15-40%) of protein-protein interactions are peptide-mediated interactions (Petsalaki and Russell, 2008), in which a short stretch of residues interact with a larger protein receptor (Mohan *et al.*, 2006). This short stretches of residues or peptide regions are often disordered alone and only obtain structure upon binding. In many cases, the peptide region is located within intrinsically disordered proteins (IDP) (Neduva, Victor *et al.*, 2005; Petsalaki and Russell, 2008) or compromise flexible linkers or loops connecting domains (Vacic *et al.*, 2007). The transient nature of these interactions makes them much harder to study experimentally compared to ordered protein-protein interactions. Thus, computational methods are crucial for guiding and designing experiments. Global peptide-docking prediction methods such as CABSdock, PIPER-FlexPepDock, pepATTRACT, and GalaxyPepDock have achieved high performances on individual benchmarks, but struggle to consistently produce reliable predictions (Kurcinski *et al.*, 2015; Alam *et al.*, 2017; Schindler *et al.*, 2015; Lee *et al.*, 2015). Local refinement predictions methods such as FlexPepDock, PEP-FOLD3, and DINC 2.0 have achieved high precision in the past, but require good starting positions (Raveh *et al.*, 2010a; Lamiable *et al.*, 2016; Antunes *et al.*, 2017). FlexPepDock for example requires a starting position within 5.5 ÅRMSD (root mean square deviation) of the correct structure to reliably produce near-native predictions (Raveh *et al.*, 2010a).

Previously we have developed InterPep (Johansson-Åkhe *et al.*, 2018) for predicting peptide-binding sites on protein surfaces. In this study, we present the improved InterPep2, which predicts the complete peptide-protein interaction complex.

## 2 Materials and Methods

### 2.1 Datasets

Two datasets of experimentally solved structures of peptide-protein interactions were used to train, evaluate, and benchmark the method in this study:

#### 2.1.1 Bound Set

The bound set consists of 502 non-redundant bound peptide-protein interaction-complexes and was previously used in InterPep (Johansson-Åkhe *et al.*, 2018). The set was randomly divided into two sets of equal sizes; a test set for benchmarking and a training set for training and tuning of the method. Validation and tuning of the method were performed on the training set by 5-fold cross-validation. The bound set is available at http://wallnerlab.org/InterPep.

#### 2.1.2 Unbound Set

In previous studies of global peptide-protein docking methods, a smaller set of 27 solved non-redundant structures of peptide-protein interaction complexes for which there exists determined unbound structures of the receptors have been used for evaluation and benchmarking of many methods (Alam *et al.*, 2017). As this set has previously been used for benchmarking, it is possible to compare against otherwise computationally heavier methods or methods which are only available through web-servers with limited programmatical access, such as PIPER-FlexPepDock (Alam *et al.*, 2017), pepATTRACT (Schindler *et al.*, 2015), and HADDOCK (Dominguez *et al.*, 2003).

As some of the targets in this set are similar to targets in the training part of the bound set, when evaluating the performance of InterPep2 on the Unbound set it was retrained for each new target, filtering training samples too similar to the target. Two complexes are considered too similar if the receptors match with a BLAST E-value of 0.05 or better.

### 2.2 Interaction Template Library

To describe possible interactions, an interaction template library was constructed from PDB (May 19, 2016) using protein-protein interactions in the defined biological units to prevent non-native interfaces from crystal packing (Carugo, O and Argos, P, 1997). The interaction template library is comprised of residues within 5.0Å (all-atoms) from another chain in a multimer complexes (intra-chain), each described by the position of its C*α* carbon.

During benchmarking, trivial templates were removed to ensure that the benchmarking measures the accuracy of the method, and not simply the difficulty of the test sets. This was done by discarding all template interfaces from complexes which receptor matched the target receptor with BLAST (Altschul *et al.*, 1997) at E-value below 0.05.

### 2.3 Performance measures

Two different criteria are used to determine if a docked conformation is successful.

- A docked peptide is “near-native” if the peptide is positioned within 4.0 Å LRMSD, i.e. the RMSD of the peptide when the complex is superimposed on the receptor is less than or equal to 4.0 Å.
- A docked peptide is considered to be at the correct binding site of the receptor if the set of receptor residues interacting with the peptide overlaps with the at least 50% of the set of receptor residues interacting with the peptide in the native complex.

Additionally, False Positive Rate (FPR) and True Positive Rate (TPR) are used to evaluate the ability to correctly rank correct and incorrect predictions:

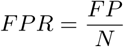

where *FP* is the number of false positives and *N* the total number of negatives in the set. FPR is also referred to as 1*−*specificity.

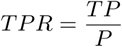

where *TP* is the number of true positives and *P* the total number of positives in the set. TPR is also referred to as recall. FPR and TPR are calculated for varying score threshold and visualized for all thresholds in a receiver operating characteristic curve (ROC-curve).

### 2.4 The InterPep2 Protocol

InterPep2 takes a protein structure and a peptide sequence as input and produces several suggested docking poses describing the peptide-protein interaction. For each of these, InterPep2 also predicts the quality of the model as measured by DockQ-score (Basu and Wallner, 2016a). The DockQ-score is a continuous measure ranging from 0.0 to 1.0, describing the overall similarity in binding between a predicted model and the native structure, based on Fnat (fraction of recovered native contacts), iRMSD (interface RMSD), and LRMSD (ligand RMSD), as by CAPRI evaluation criteria (Basu and Wallner, 2016a).

The complete protocol consists of five steps, explained in detail below:

#### 2.4.1 Template Alignments

Every full chain with one or more interfaces included in the Interaction Template Library is aligned to the receptor protein structure using TM-Align (Zhang and Skolnick, 2005). For each alignment, the interfaces of the aligned chain are superimposed on the receptor by the same rotational matrix used for the full chain in the TM-Align alignment, creating templates for interaction.

#### 2.4.2 Peptide Conformations

The peptide conformational space is represented by 50 structural models of the peptide generated using structural fragments from known structures with similar sequence and secondary structure. In detail, PSI-BLAST (Altschul *et al.*, 1997) is used to generate a sequence profile for the peptide, which is used in PSI-PRED (Jones, 1999) to predict secondary structure. The sequence profile and the secondary structure prediction are both used in the Rosetta Fragment Picker application (Gront *et al.*, 2011), which selects 50 fragments from 2000 decoys by finding sequence and secondary structure matches in a representative set of monomeric protein structures. The 50 fragments are then extracted from their full structures, and sequence is changed to that of the query peptide using the Rosetta fixbb application (Kuhlman and Baker, 2000). This protocol for generating peptide fragments is similar to that of PIPER-FlexPepDock (Alam *et al.*, 2017), and has been shown to reliably sample near-native peptide conformations.

#### 2.4.3 Build interaction complex

Interaction complexes are constructed by combining the 2,500 best template alignments (by TM-score) from the template alignment step with the 50 conformations of the peptide, resulting in 125,000 (2,500 x 50) coarse interaction models. The models are created by, for each combination of receptor-to-interaction-template alignment and peptide conformation, aligning the peptide to the complementary side of the template interface using InterComp (Mirabello and Wallner, 2018). InterComp is used in favor of TM-Align as it can align with respect only to coordinates and amino acid identity, completely eschewing sequence order (Mirabello and Wallner, 2018). This means that even a composite interface formed from non-consecutive residues in any order can align to a short straight peptide.

**Figure 1:**
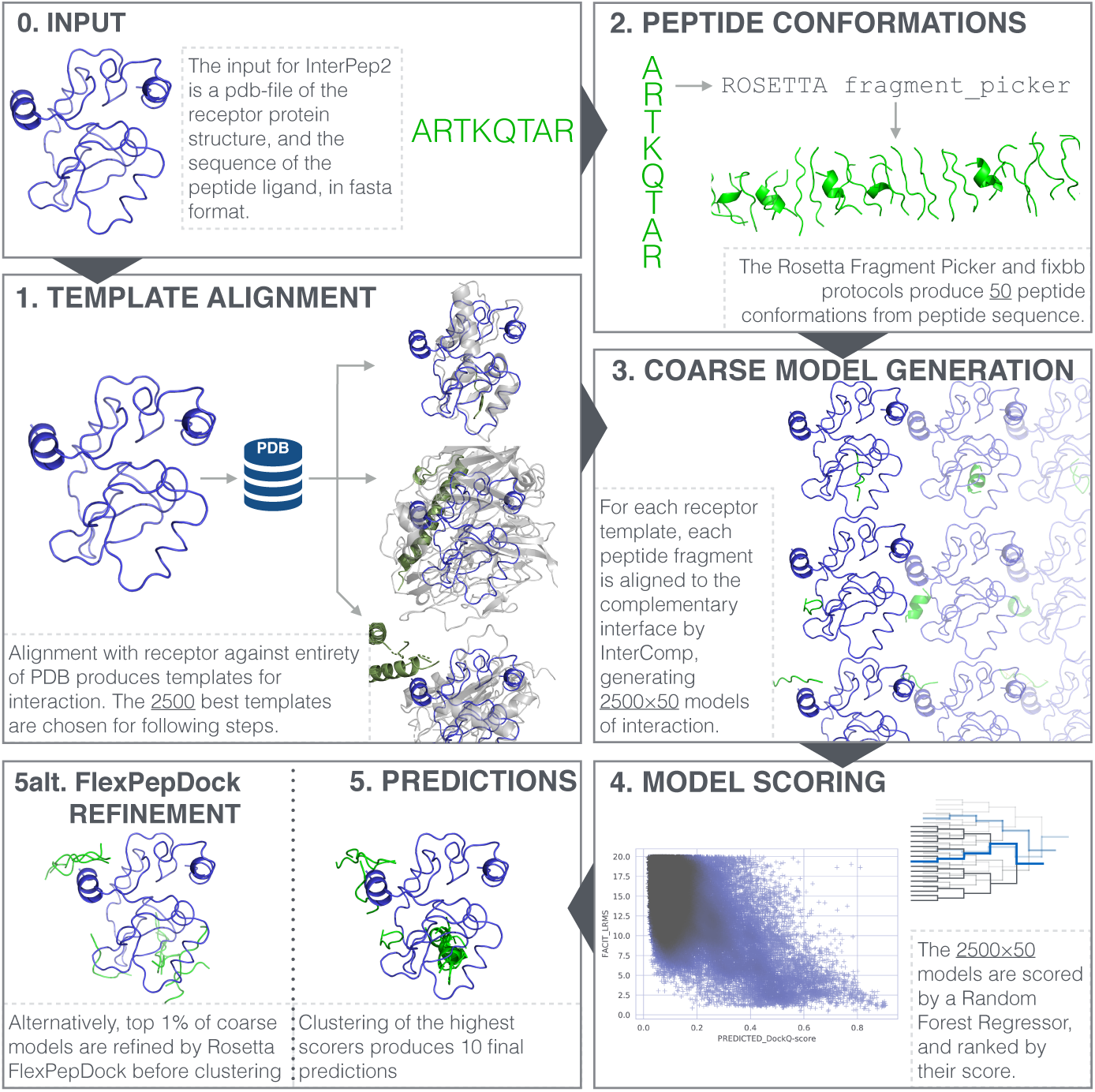
Summary of the InterPep2 method. The receptor used in the example is of the DNMT3a ADD domain, and the peptide is a H3 peptide (pdb-id 4QBQ for original complex (Noh *et al.*, 2015)).

#### 2.4.4 Model Scoring

Each suggested model of the interaction complex is scored by a regression random forest trained to predict the DockQ-score of the complex, based on a number of features relevant to peptide-protein interactions (Table 1). During the development of InterPep2, additional features were considered, but only features, which showed considerable contribution to performance were kept. The parameters of the random forest as well as the selection of features was optimized by 5-fold cross-validation on the training data, details in Supplementary Information.

**Table 1:**
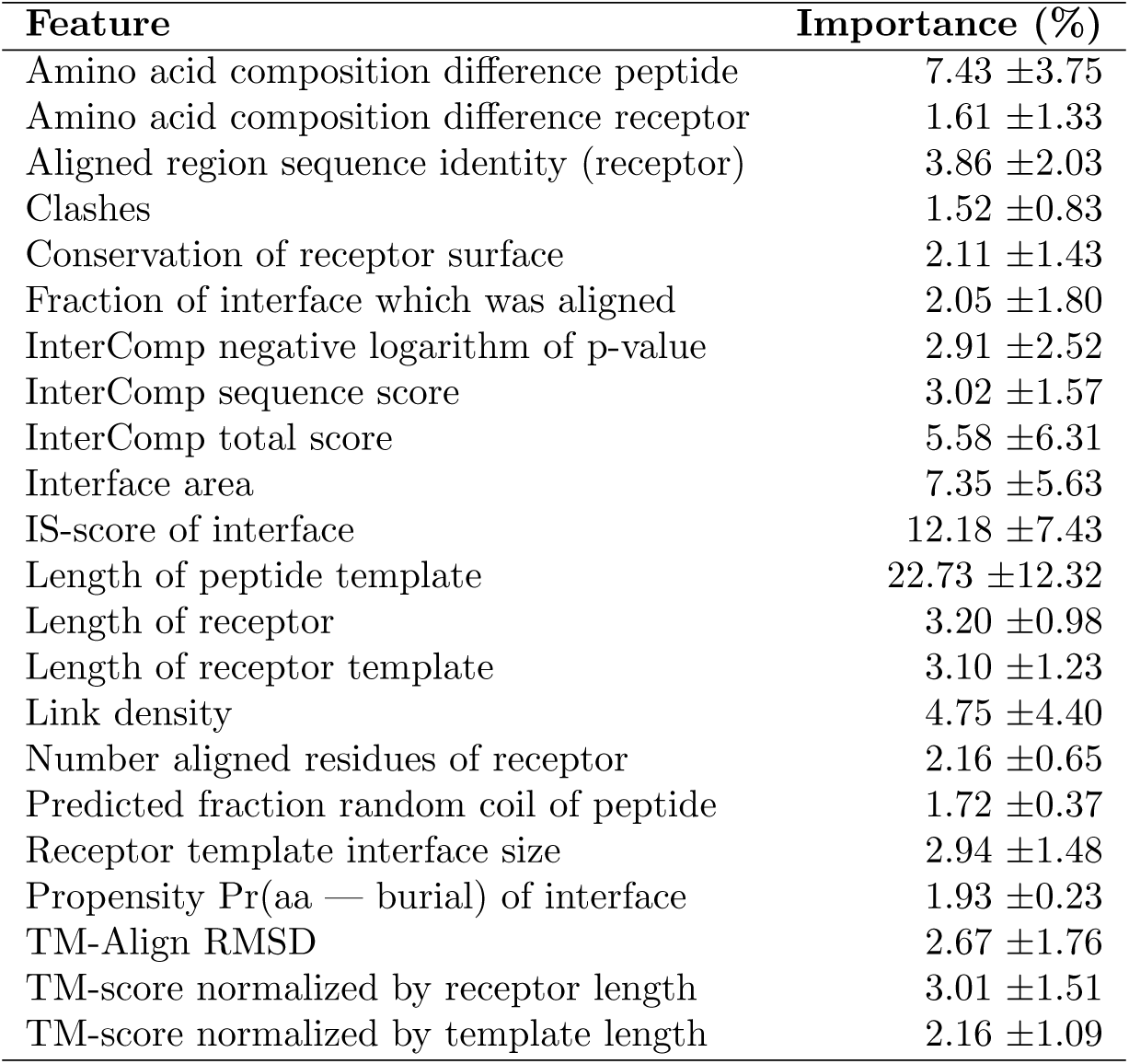
Table of features used by the Random Forest, and their relative importance as measured by reduction in Gini Impurity. Note that the features regarding TM-Align refers to the alignment between the receptor and its template, and the features regarding InterComp refers to the alignment between the peptide and its template.

The random forest of InterPep2 is implemented using the Scikit-learn python package (Pedregosa *et al.*, 2011).

#### 2.4.5 Predictions

The final predictions are ranked by the predicted DockQ-score and filtered to ensure no two predictions are within 4.0 Å LRMSD.

#### 2.4.6 Refinement (optional last step)

InterPep2 generates unrefined coarse interaction models that could benefit from refinement. To analyze the usefulness of refinement, the option to refine the coarse models using the Rosetta FlexPepDock refinement protocol (Raveh *et al.*, 2010b) was added (InterPep2-Refined). Using this option, up to 12,500 of the best ranking predictions with a predicted DockQ-score above 0.2 are refined, and the top 1% (top 125) models with best Rosetta reweighted scores are clustered at 2 Å LRMSD, the clusters are ranked by the mean predicted DockQ-score of the models in the cluster, and the cluster centers are given as final predictions.

## 3 Results and Discussion

### 3.1 Database Coverage

To get an estimate on the upper bound of prediction performance, the available templates for the 251 complexes in the bound set were analyzed. Using different subsets of the available templates, the performance was assessed on three levels: high quality (LRMSD*≤*4.0Å), at least medium quality (LRMSD*≤*5.5Å), and at least correct site on the receptor, see Table 2. In all cases, templates from closely related structures are removed (BLAST e¡0.05).

**Table 2:** Coverage of templates for the 251 complexes in the bound set, i.e. the upper bound on the performance. ≤*4.0Å* and ≤*5.5Å* describe for how many complexes that can be modeled within 4.0 or 5.5 Å LRMSD, respectively. *Correct Site* refers to how many complexes have at least one template that positions the peptide in the correct site on the receptor.

| Template set | $\leq 4.0\text{\AA}$ | $\leq 5.5\text{\AA}$ | Correct site |
| --- | --- | --- | --- |
| All inter-chain templates | 89 | 164 | 194 |
| Peptide-protein templates | 70 | 109 | 140 |
| Protein-protein templates | 59 | 136 | 168 |

Looking at Table 2, we can see that by using only peptide-protein templates, it is possible to model 70 of the 251 test complexes at high quality, while adding protein-protein templates increases this number to 89/251. In 22 of these 89 cases (24%), it is optimal to use a protein-protein rather than a peptide-protein interaction as a basis for the template, although it is possible to produce near-native models for 59 targets using only protein-protein templates. Given that a considerable fraction of the high-quality models originate from protein-protein templates, there is a substantial gain by extending the database to cover both protein-protein and peptide-protein templates, and not limiting the search-space to peptide-protein interactions only.

Notably, there are more high-quality templates from peptide-protein templates compared to protein-protein templates, while the opposite is true for medium quality or correct site templates. This implies that peptide-protein templates are generally better templates for other peptide-protein interactions, while it is still viable to predict peptide-protein interactions with protein-protein templates. The fact that there are far more protein-protein templates than peptide-protein templates in the template set certainly increase the chances of finding potential useful templates.

### 3.2 Benchmark: Bound Set

The performance of InterPep2 and InterPep2-Refined was compared to two other available established methods: GalaxyPepDock (Lee *et al.*, 2015) and CABSdock (Kurcinski *et al.*, 2015) on the bound set. Many other methods for global peptide to protein receptor docking exist, but most are not readily available to be run in large-scale tests, having either no standalone version or are only available through web-servers with limited programmatical access. Since InterPep2, InterPep2-Refined, and GalaxyPepDock are template-based, trivial templates (BLAST e¡0.05) were not allowed so as to emulate a real-world situation.

First the ability of each method to accurately model the proteins in the bound set at three quality levels, near-native at top1, near-native among top10, and correct site modeling at top1, were assessed (Figure 2). InterPep2 has more near-native models at top1 compared to GalaxyPepDock, and CABSdock, it also has more models with the peptide at the correct site; and applying FlexPepDock refinement in InterPep2-Refined starting from InterPep2 models consistently improves the results.

**Figure 2:**
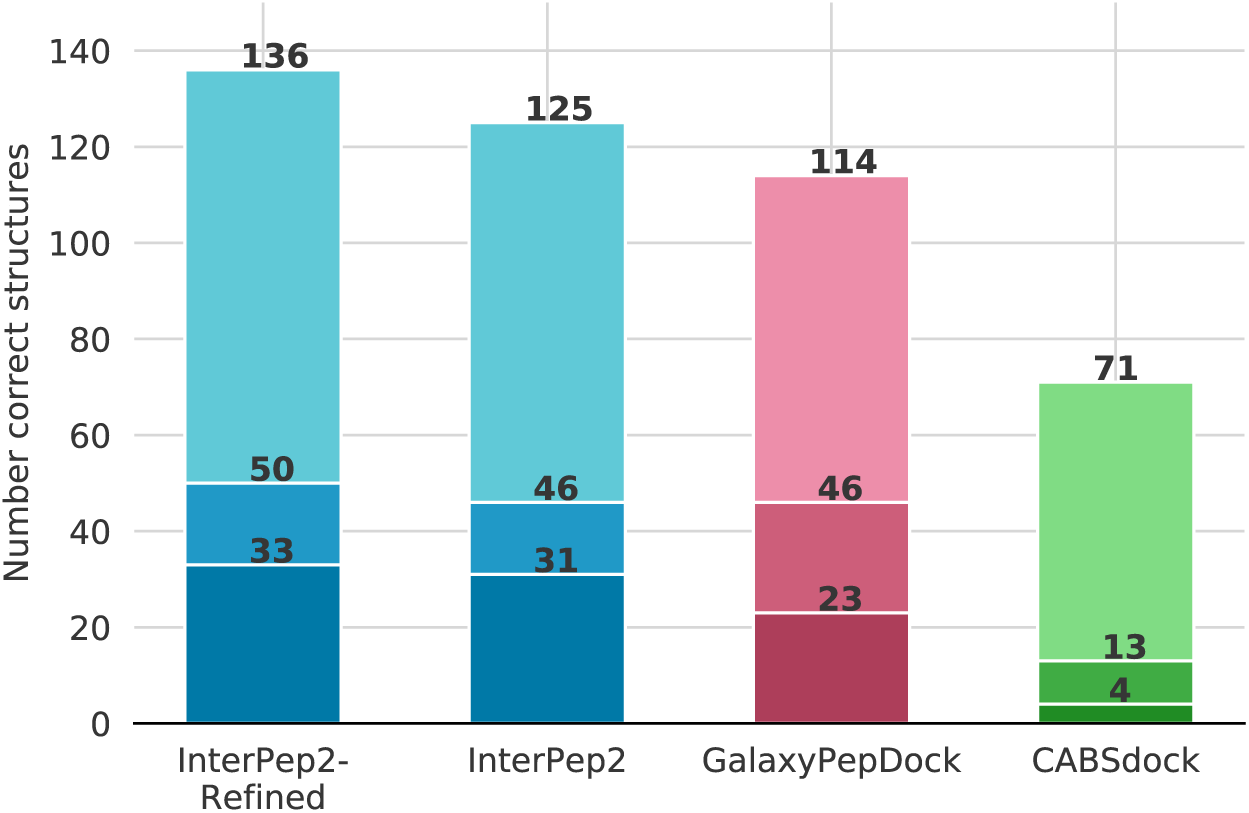
Number of correct model structures for the different methods for varying quality threshold. From bottom to top (dark to lighter shades), number of near-native models (LRMSD≤4.0Å) at top1, number of near-native models choosing the best of top10 for each target, and number of top1 models with the peptide at the correct site.

Both InterPep2 and GalaxyPepDock provide model scores, InterPep2 in the form of the predicted DockQ-score and GalaxyPepDock as a predicted accuracy. The ability of these scores to separate near-native from non near-native predictions were investigated in a ROC-curve (Figure 3). Shown here, InterPep2 also has a larger AUC and recalls more true positives across all specificity levels.

**Figure 3:**
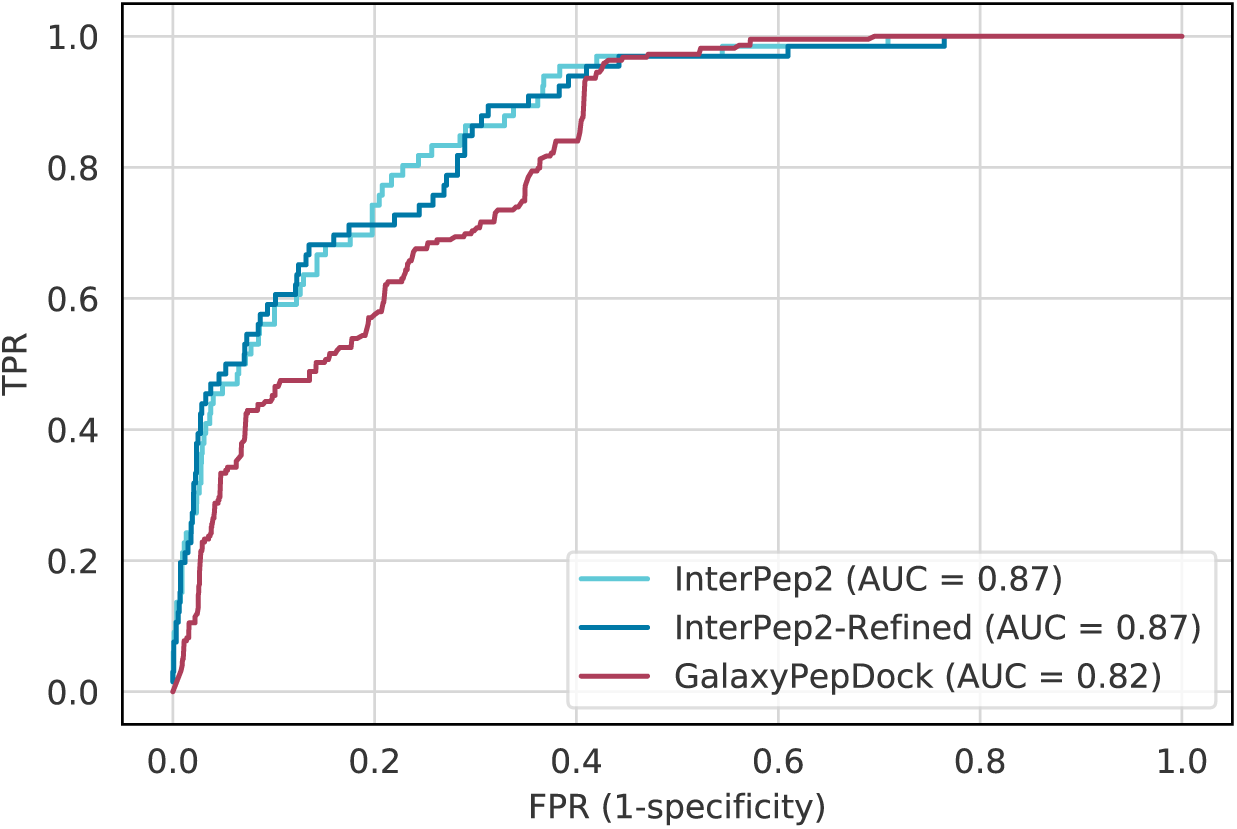
Receiver Operating Characteristic (ROC) curve showing the capability of InterPep2 and GalaxyPepDock to correctly rank their own predictions.

Additionally, to test the initial hypothesis that templates from structured proteins can be used to predict the binding of regions originating from natively disordered proteins, peptides in the bound set was classified as disordered or ordered. A peptide was considered disordered if it matched an annotated disordered region from DISPROT (Piovesan *et al.*, 2016) with a BLAST E-value of 0.05 or less, otherwise it was classified as ordered. The difference in loss in DockQ-score when predicting complexes including disordered and ordered peptides was analyzed (Supplementary Figure S2). Based on Kolmogorov-Smirnov and two-sided t-tests it could be concluded that there was no difference between predictions made for disordered compared to ordered peptides (P*>*0.5 for the null hypothesis that there is a difference).

### 3.3 Benchmark: Unbound Set

Note that when benchmarking against the unbound set, the template-based methods are still not allowed to use templates closely related to the target receptors. As can be seen form Figure 4, most methods perform similarly to each other. However, PIPER-FlexPepDock produces the most near-native structures when looking at top10: 18/27, compared to InterPep2-Refined at 15/27.

**Figure 4:**
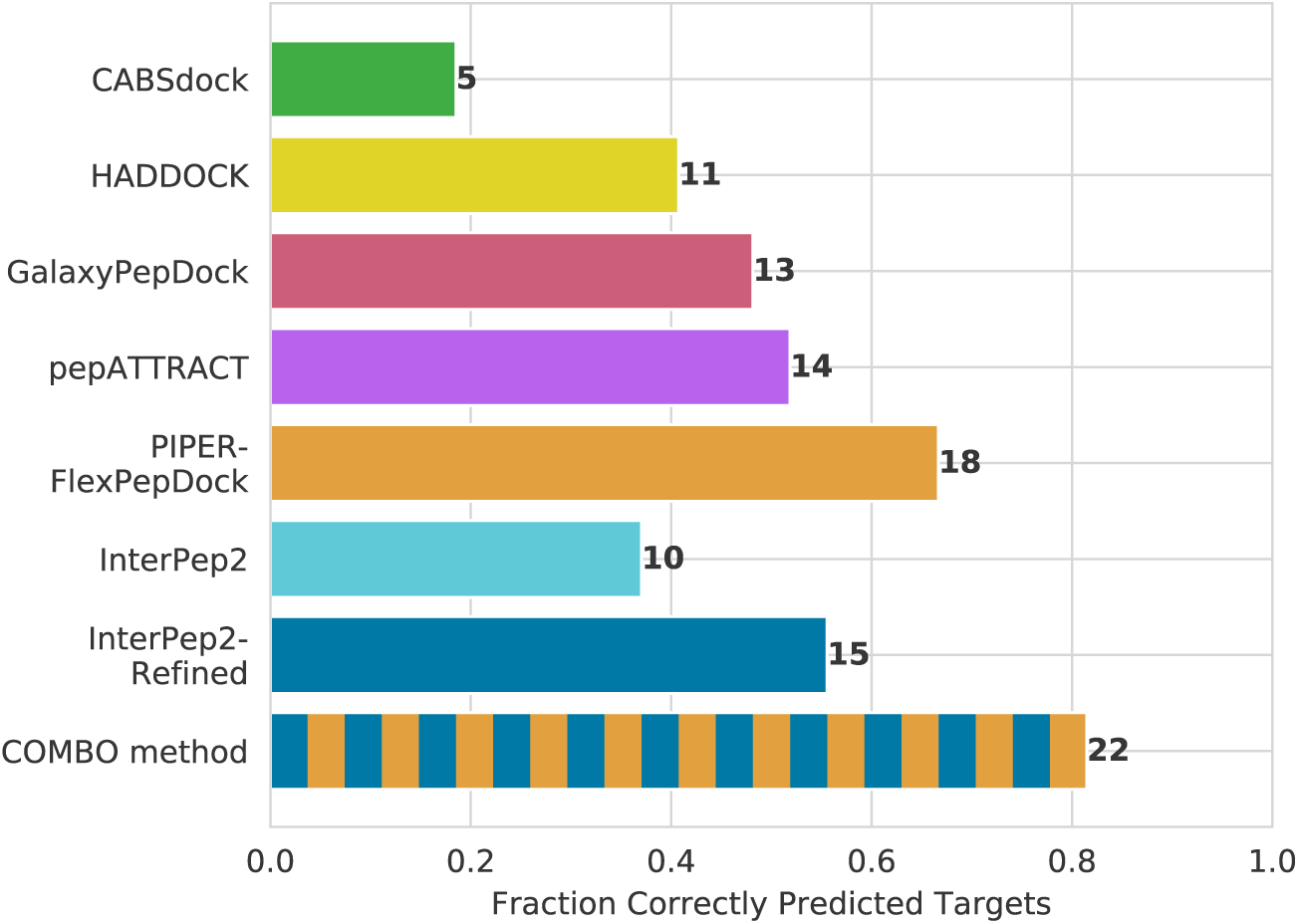
Comparison of the ability of different methods to produce near-native docked peptide structures on the unbound set. As previously, a prediction counts as correct if within 4.0 Å LRMSD from the native. Data for CABSdock, HADDOCK, pepATTRACT, and PIPER-FlexPepDock is taken directly from (Alam *et al.*, 2017), in the form of near-native among top10, which is why no data regarding top1 or how many peptides are positioned at the correct site is shown.

#### 3.3.1 Combo method

Even though PIPER-FlexPepDock and InterPep2-Refined produce the highest numbers of near-native structures, the overlap between the predictions from these methods is relatively small: 6 structures correctly modeled by InterPep2-Refined are not correctly modeled by PIPER-FlexPepDock (see Supplementary Figure S4). It is trivial to combine these methods into a combination method superior to both alone, denoted *Combo method* in which InterPep2-Refined models are selected if the InterPep2 predicted DockQ-score is above a threshold X (X=0.412*±*0.009, optimized using leave-one-out cross-validation see Supplementary Information), otherwise a PIPER-FlexPepDock prediction is selected.

#### 3.3.2 Impact of Refinement

As seen in Figure 2 and 4, there is an inconsistency in how InterPep2 performs in relation to CABS-dock and GalaxyPepDock, with InterPep2 performing worse than GalaxyPepDock in the unbound test but better in the bound test. Most likely, this inconsistency stems from the fact that InterPep2 is only trained on bound structures, leading to performance loss when predicting bound structures starting from unbound. Additionally, the relative difference in performance between InterPep2 and InterPep2-Refined is much larger when the method is applied to the unbound set, indicating that the refinement can rescue more targets for the unbound compared to the bound set, probably partially because of allowing small backbone changes. Finally, the larger bound set contains peptides in the ranges of 5 to 25 residues long, which is much more difficult to predict than the 5 to 15 residues long peptides of the unbound set, accounting for the difference in fraction successful predictions for InterPep2-Refined (55.6% near-native predictions for the unbound set with only 19.9% near-native predictions for the bound set).

### 3.4 Features of a Good Template

As shown in Figures 2 and 4, the random forest can perform well on data not initially trained on, and can therefore be assumed generalizable. As such, the features of high importance in Table 1 should represent generally important features for the similarity of one peptide-protein interaction interface to another inter-chain interface. Four features stand out as the most important:

The feature with highest importance is the length of the chain representing the peptide in the template. This is unsurprising, as we have previously concluded that for acquiring near-native predictions it is often better to use peptide-protein interactions as templates, rather than protein-protein interactions (see Table 2 and Figure S3). Among all predictions made by InterPep2 on the bound set, 179 of the 251 (71.3%) top ranking predictions are derived from peptide-protein template interactions. The second-most important feature is the IS-score of the suggested interface, representing the conservation of the interaction surface. Sequence conservation is used as part of many machine-learning-based approaches to modeling and can sometimes alone be enough for the identification of binding-sites (Mayrose *et al.*, 2004). Finally, the similarity in amino acid composition between the peptide and its template, and the total surface area covered by the peptide, also have a large importance. In summary, features that describe the similarity of the peptide to its template, and features that identify the sequence conservation and size of the binding area of the receptor seem to be the most important.

More detailed features such as the residue packing or shape complementarity that previously have been proven important when ranking refined protein-protein models (Basu and Wallner, 2016b), do not impact the result at all when ranking unrefined coarse models as is the case here (see Supplementary Information). The number of clashes do have an effect on prediction accuracy, but the importance is small (Table 1).

### 3.5 Prediction Example

An example of a successful InterPep2 prediction is shown in Figure 5. This example is of the ADD domain binding to a H3 peptide tail. Note how in Figure 5A, predictions 7-10 suggest an alternate binding site. The templates which suggest this site are all from the polycomb protein EED, which has also shown to bind to the H3 peptide in previous studies (Li *et al.*, 2014). Additionally, analyzing the general areas of interaction of all predictions with ConSurf (Ashkenazy *et al.*, 2016; Landau *et al.*, 2005) showed both binding sites to be considerably more conserved than the rest of the protein surface. Both predicted binding sites had relative evolutionary rate scores with a mean of −0.56, implying conservation, compared to −0.89 for the center of the receptor (heavily conserved), and contrasting the mean of 0.87 on the rest of the surface (little to no conservation), indicating a possible alternate interaction-site. In another structure of the domain, 3QL9, the protein is shown with its C-terminal helix in this groove, and in 2PVN this site is where the ADD domain interfaces with the other domains of the full DNMT3L structure.

**Figure 5:**
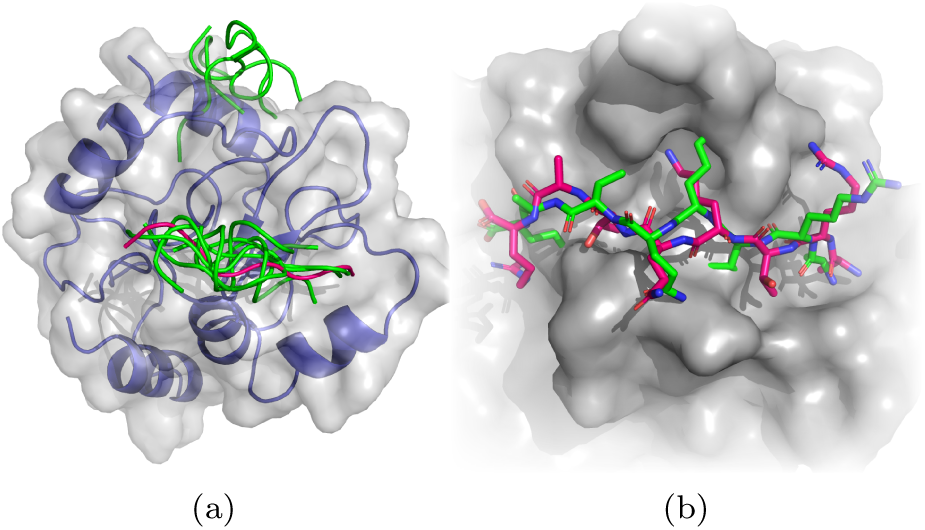
An example of a successful InterPep2 prediction on the complex of DNMT3a ADD domain binding to H3 peptide (PDB: 4QBQ (Noh *et al.*, 2015)). **A)** The top 10 predictions made by InterPep2, peptides in green, as well as the native structure, peptide in pink. The receptor is blue and its surface a semitransparent gray. **B)** A closer look at the top 1 prediction from InterPep, in green, together with the native peptide, in pink. The predicted peptide is positioned 2.1 Å RMSD from the native peptide conformation, counting backbone positions. The images were constructed through PyMOL (Schrödinger, LLC, 2015).

This example shows that even though some of the top predictions might not be at the correct peptide binding site, they might indicate other areas of biological importance. To test this hypothesis, we analyzed all top 10 predictions for all test targets (2510 predictions in total), for which a total of 1614 predictions position the peptide at the correct binding sites. By comparing to binding sites of closely related structures within the PDB (BLAST e¡1e^*−*20^), 401 of the remaining 896 predictions position the peptide at another binding site, which is either for another peptide or another protein-protein interaction, sometimes dimerization. A histogram over the distributions of the predicted score for the different types of predicted sites, Figure 6, shows that, although nonsense-predictions and predictions of other sites (sites which are not the peptide-protein site we test for) are difficult to tell apart, they follow different distributions. In general, at a low predicted score of circa 0.15 to 0.25, it is roughly equally likely that the prediction is of the correct site as that it would be of another binding site or a nonsense-predictions altogether, while at higher predicted scores the majority of predictions are clearly most often positioned at the correct binding site.

**Figure 6:**
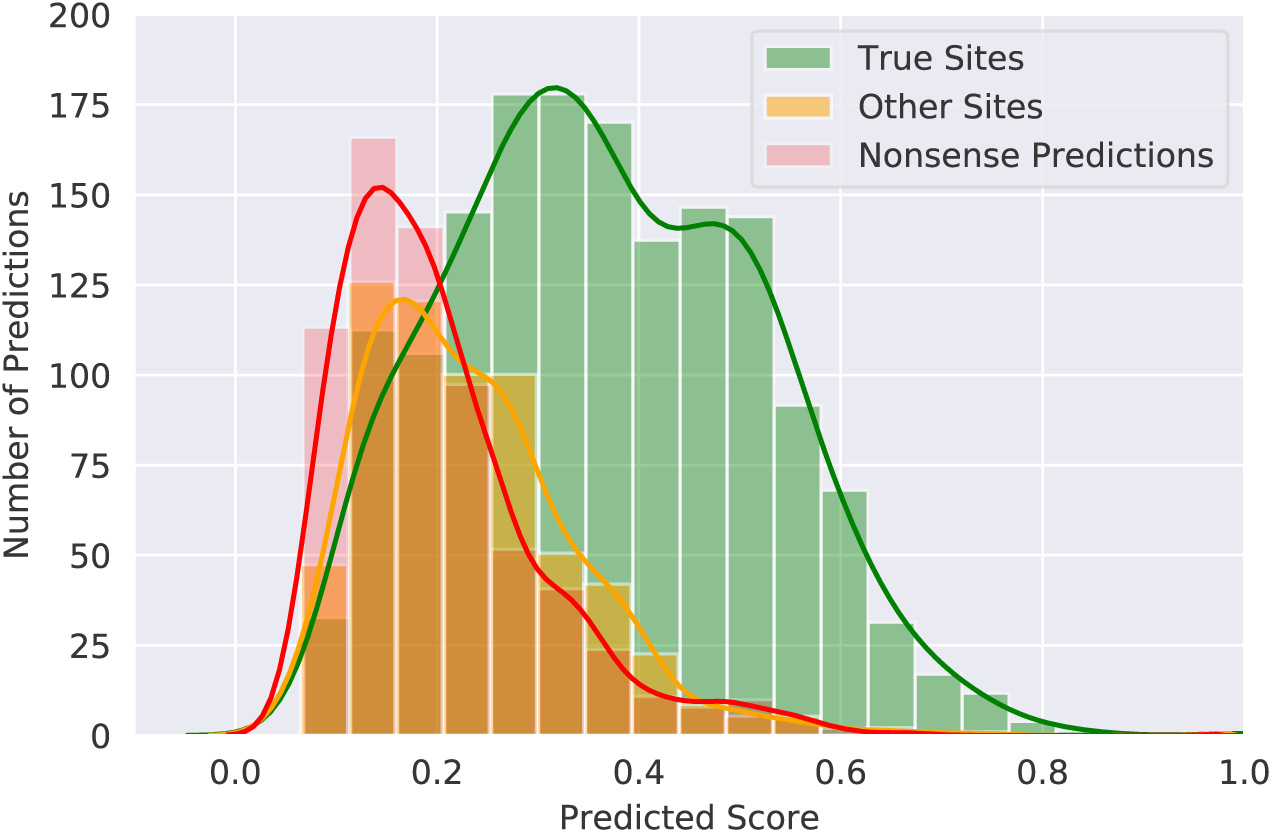
Histogram for the predicted DockQ scores of the top 10 predictions for every target, separated by if the prediction found the correct site, another kind of binding site, or an incorrect site. All three distributions are significantly different, with P*<*0.0001 using d by two-sided Kolmogorov-Smirnov-test.

## 4 Conclusions

InterPep2 applies structural templates for docking peptide fragments, achieving greater accuracy than most other state-of-the-art methods, even when limiting the training-data and template-database to structures without significant sequence similarity to the targets predicted. On a difficult data-set of bound interactions presented here, InterPep2-Refined successfully places 33 of 251 peptides within 4.0 Å of their native positions, with a True Positive Rate of 59% at a False Positive Rate of 10%. Even in cases where the peptide was not placed within 4.0 Å LRMS, it was placed at the correct binding-site a majority of the times; 136 out of 251.

On a frequently used dataset of 27 unbound-to-bound complexes, InterPep2Refined performed second-best, successfully predicting 15 of 27 ligand conformations. More interesting however, is the prospect of combining InterPep2-Refined with PIPER-FlexPepDock by simply running InterPep2 first and continuing with either InterPep2-Refined or PIPER-FlexPepDock depending on the InterPep2-score. This combined method vastly outperformed both methods it was derived from, successfully generating near-native models for 22 of the 27 conformations.

## Acknowledgments

This work was supported by a Swedish Research Council grant, 2016-05369, The Swedish e-Science Research Center, and the Foundation Blanceflor Boncompagni Ludovisi, née Bildt. The computations were performed on resources provided by the Swedish National Infrastructure for Computing (SNIC) at the National Supercomputer Centre (NSC) in Linköping.

