## Supplementary Information for "InterPep2: Global Peptide-Protein Docking with Structural Templates"

### InterPep2 Supplementary Information

October 21, 2019

#### Random Forest Features

##### Final Features

Following are descriptions of all features used by the random forest in the final version of InterPep2.

###### Amino Acid Composition Difference Peptide & Amino Acid Composition Difference Receptor

These features were calculated as in Johansson-Åkhe *et al.* (2018): For both of the elements compared (receptor interaction-surface to receptor template interaction-surface or peptide to peptide template), a composition vector of 20 values is calculated, where each value represents the percentile propensity of a certain amino acid in the sequence. This vector is then multiplied by the BLOSUM62 matrix (as shown in Equation 1), and the angle between the new vectors (after multiplication with the matrix) is the Amino Acid Composition Difference (Equation 2).

$$\vec{y}_i = B\vec{x}_i \tag{1}$$

where $\vec{x}_i$ is the amino acid composition vector of sequence $i$, and $B$ is the BLOSUM62 matrix.

$$\theta_{jk} = \arccos\left(\frac{\vec{y}_j \cdot \vec{y}_k}{|\vec{y}_j| + |\vec{y}_k|}\right) \tag{2}$$

where $\theta_{jk}$ is the Amino Acid Composition Difference between sequences $j$ and $k$.

###### Aligned Region Sequence Identity (receptor)

The sequence identity between the receptor and receptor template in the region successfully aligned by TMalign Zhang and Skolnick (2005).

##### Clashes

The number of clashes between the positioned peptide and the receptor. A clash is defined as the closest pair of heavy atoms between residues being closer than 2.2 Å.

##### Conservation of Receptor Surface

The relative conservation of the residues in contact with the positioned peptide, divided by the relative conservation of the rest of the receptor surface. For this purpose, a residue is defined as a surface residue if at least 15% exposed. The conservation is calculated using PSIBLAST Altschul *et al.* (1997).

##### Fraction of Interface which was Aligned

The fraction of interacting residues in the receptor which were successfully aligned to the receptor template using TMalign.

##### InterComp Negative Logarithm of p-value

The negative logarithm of the p-value output from InterComp Mirabello and Wallner (2018) aligning the peptide to the peptide template surface.

##### InterComp Sequence Score

The sequence similarity score output from InterComp aligning the peptide to the peptide template surface.

##### InterComp Total Score

The final total score output from InterComp aligning the peptide to the peptide template surface, which is a combination of sequence and structural similarity.

##### Interface Area

The size of the area of contact between the receptor and positioned peptide, as calculated by NIP-NSc Mitra and Pal (2010).

##### IS-score

The IS-score of the receptor residues in contact with the positioned peptide, calculated as in Bertoni *et al.* (2017) with a 40% sequence identity inclusion threshold:

First, the relative entropy $RE$ is calculated for each column $c$ of a multiple sequence alignment acquired from HHBLITS Alva *et al.* (2016).

$$RE_c = \sum^{a} p_a \log_2 \frac{p_a}{p_{ab}} \tag{3}$$

where $p_a$ is the probability of an amino acid $a$ to be in column $c$, and $p_{ab}$ is the background distribution of $a$ in the entire alignment.

Next, for both the interacting surface and the rest of the protein, a weighted average of entropy $\langle S \rangle$ is calculated ($\langle S \rangle_i$ for the interacting surface and $\langle S \rangle_s$ for the rest of the protein).

$$\langle S \rangle = \frac{\sum rASA_c RE_c}{\sum rASA_c} \tag{4}$$

where $rASA_c$ is the relative solvent accessible surface area of residue $c$.

Finally, the IS-score is defined as a ratio between the two:

$$IS = \ln \frac{1 + \langle S \rangle_s}{1 + \langle S \rangle_i} \tag{5}$$

##### Length of Peptide Template

The number of residues in the full chain of the peptide template (not only the interacting part).

##### Length of Receptor

The number of residues in the receptor structure.

##### Length of Receptor Template

The number of residues in the receptor template (not just the interacting part).

##### Link Density

The number of residue pairs between the receptor and peptide positioned within 6.0 Å of each other, divided by the length of the peptide. Note that one residue can be included in several pairs.

##### Number of Aligned Residues of Receptor

The number of receptor residues successfully aligned by TMalign when aligning against the receptor template.

##### Predicted Fraction Random Coil of Peptide

The fraction of the peptide sequence predicted to be random coil by PSI-PRED Jones (1999).

##### Receptor Template Interface Size

The number of residues in the receptor template's interface.

##### Statisics P(aa | burial) of the interface

Amino acid propensity given the burial state of the amino acid, calculated same as residue given burial (rGb) in Basu *et al.* (2014):

$$P(aa|burial) = \frac{1}{N_{res}} \sum_{i=1}^{N_{res}} \log_{10}(Pr_i) \tag{6}$$

where $Pr_i$ is the propensity of the amino acid at position $i$ for the particular degree of relative exposure it has in this position. Exposure is divided into 4 bins with divisions at 0.05, 0.15, and 0.3 fraction relative exposure, as compared to an Ala-X-Ala extended conformation.

##### TMalign RMSD

The RMSD reported by TMalign after aligning the receptor to the receptor template.

##### TMscore normalized by receptor length

The TMscore reported by TMalign after aligning the receptor to the receptor template, as normalized by receptor length.

##### TMscore normalized by template length

The TMscore reported by TMalign after aligning the receptor to the receptor template, as normalized by receptor template length.

#### Filtered features

These features were initially also included, but were filtered from the final method as they had the lowest mean feature importances or highest importance variation to mean importance ratios, and removing hem had negligible or no effect on cross-validation performance.

##### Domain-domain interface

One-hot encoding describing if the template of interaction was from an intra-chain interaction rather than an inter-chain interaction.

##### InterComp p-value

The p-value output from InterComp aligning the peptide to the peptide template surface.

##### Interface Contact Preference

The interface contact preference score is calculated by summing the normalized residue-residue contact scores of Glaser *et al.* (2001) for each pair of receptor-peptide interacting residues and dividing by the total number of contacts.

##### Interface Packing

Shape complementarity as calculated by NSc Mitra and Pal (2010).

##### Length of Peptide

The number of residues in the peptide sequence.

##### Mean Sequential Distance of Peptide Template Interface

As calculated by the following:

$$MSD = \frac{\sum_{i \in k} s_i - s_{i-1}}{N} \tag{7}$$

where $k$ is a list of the residues in the template interface sorted by sequence position, $s_i$ is the sequence position of residue $i$, and $N$ is the total number of residues in $k$.

##### Physical Dimensions of the Peptide Template Interface

The principal components of inertia of an ellipsoid with the same moments of inertia as the peptide, given as length, width, and height of the ellipsoid. Calculated as in Wallner and Elofsson (2003).

##### Peptide Posttranslational Modification

Three dummy variables of 0.0 or 1.0 described whether the peptide was acetylated, methylated, or contained some other posttranslational modification.

##### Predicted fraction $\alpha$-helix and $\beta$-sheet of peptide

As predicted by PSI-PRED Jones (1999).

##### Secondary Structure Composition Peptide Template

The fraction $\alpha$-helix, $\beta$-sheet, and coil of the interaction-surface of the peptide template, as calculated using STRIDE Frishman and Argos (1995).

##### Shape Complementarity

The shape complementarity of the interface between the receptor and positioned peptide, as calculated using NIP Mitra and Pal (2010).

| **Parameter** | **Range** |
|---|---|
| Maximum depth | 10 to 150, None |
| Max features considered per split | N/3, N/2, 2N/3, auto, $\sqrt{N}$ |
| Minimum amount of samples per leaf | 1 to 1000 |
| Minimum amount of samples per split | 1 to 1000 |
| Number of trees | 100 to 3000 |

Table S1: Search-space for random forest parameters. In the "max features considered per split" case, "auto" sets to $\sqrt{N}$ when using a classifier random forest, and N when using a regressor random forest.

#### Random Forest Optimization

The final set of features was optimized from the expanded set by iteratively dropping the features with the least importance as measured by Gini Impurity until significant negative effects in loss or correlation to true appeared. This was done with 5-fold cross-validation on the training-set.

After filtering features by importance, the parameters of the random forest were optimized by a random grid-search, investigating several ranges of parameters as shown in table S1.

After the parameter optimization, the iterative filtering of features by importance restarted with all features. Lastly, features were iteratively filtered by variance in importance to importance ratio, with the otherwise same criteria for removing features as for the previous feature filtering steps.

##### Normalization

To discourage the random forest from focusing on predicting whether a given receptor target is easy or difficult, and focus on ranking the different peptide conformations. It was investigated if normalizing the training data for each target would improve ranking. There proved to be significant difference in favor of normalization when evaluating on overall correlation between predicted DockQ-score of complexes and true DockQ-scores (p-value 0.047, wilcoxon signed rank test, "true" DockQ-score being the normalized DockQ-score in this case). However, since the ultimate goal is to find the best template, the loss in DockQ-score between the actual best complex and the complex predicted as best by the random forest should more accurately describe the performance of the method. In this case, there was no significant difference between normalizing and not normalizing training data (p-value 0.74), especially not when disregarding obvious templates (p-value 0.93).

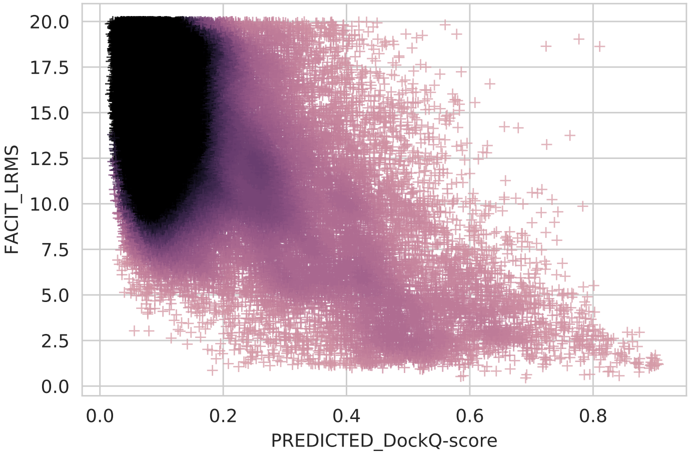


Figure S1: Distribution of LRMSD of different models versus their predicted DockQ-scores, as predicted by the random forest. Note the clearly lowered concentration of sub-4 Å structures at less than 0.2 predicted DockQ-score. The Figure has been cropped not to show results over 20.0 Å LRMSD. For ease of comparison and view, only 250,000 randomly selected models are shown.

### FlexPepDock Refinement

The FlexPepDock refinement step is time-consuming. To avoid refining hopeless cases, where the starting structures are very far from the native, the ability of the predicted DockQ-score to estimate the actual LRMSD was analyzed (Figure S1). For predicted DockQ-score below 0.2 very few starting structure are below 4 Å. This shifts at predicted DockQ-score above 0.4, where the vast majority of the structures instead are below 4 Å. This is in agreement with the observed increase in correctly identified binding sites at 0.4 predicted DockQ-score reported in the main text. Based on the above, it was decided that only starting structure with a predicted DockQ-score above 0.2 should be refined.

### Peptide fragments from disordered proteins

50,000 datapoints were randomly sampled from predictions of complexes involving peptide fragments from disordered proteins and ordered peptides, respectively (Figure S2).

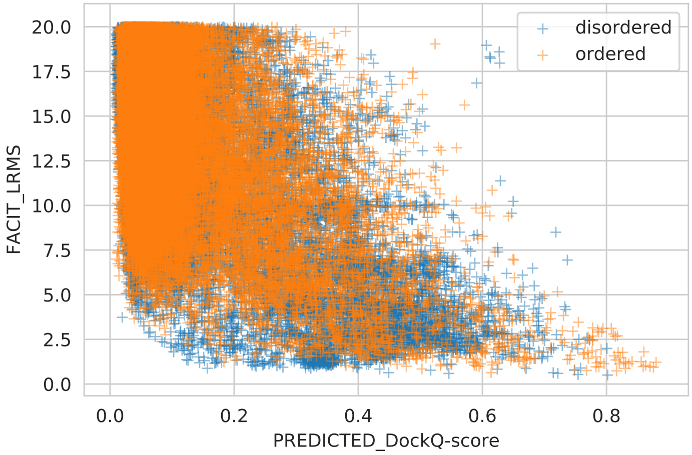


Figure S2: Predicted DockQ-scores vs. LRMSD for disordered and ordered peptides.

### Domain-Domain Interfaces

In addition to using protein-protein and peptide-protein interfaces, domain-domain interfaces were also investigated as potential templates, as previous studies have shown that the bonds in such interfaces are similar to those between proteins and peptides Vanhee *et al.* (2009).

Including domain-domain interfaces in the Interaction Template Library increased the total size of the library by 57% (from 561,936 to 880,956 interfaces), but the cross-validated performance difference was not significant (p-value $> 0.1$).

Table S2 shows the increase in possible corrects when including domain-domain interfaces as templates. Although there is a difference, the increase is very small considering the amount of additional possible interfaces, which results in an imbalanced set, where, because of the likelihood of a false positive, the Random Forest is better off always choosing protein-protein or peptide-protein interfaces as templates rather than domain-domain interfaces, see Figure S3.

### InterPep2-PIPER-FlexPepDock (combo)

In Figure S4, the complementary difference in predictions between InterPep2-Refined and PIPER-FlexPepDock Alam *et al.* (2017) is shown. The predictions from the two methods are quite different, but more importantly for a majority of the cases at least one method have near-native prediction below 4.0 Å LRMSD. This fact is used by the Combo method, which uses an automatically selected

| **Template set** | **≤4.0Å** | **≤5.5Å** | **Correct site** |
|---|---|---|---|
| All templates | 99 | 181 | 212 |
| All inter-chain templates | 89 | 164 | 194 |
| Peptide-protein templates | 70 | 109 | 140 |
| Protein-protein templates | 59 | 136 | 168 |
| Domain-domain templates | 30 | 106 | 170 |

Table S2: Coverage of templates for the 251 complexes in the test set, i.e. the upper bound on the performance. Extended to include domain-domain templates. *≤4.0Å* and *≤5.5Å* describe for how many complexes that can be modeled within 4.0 or 5.5 Å LRMSD, respectively. *Correct Site* refers to how many complexes have at least one template that positions the peptide in the correct site on the receptor.

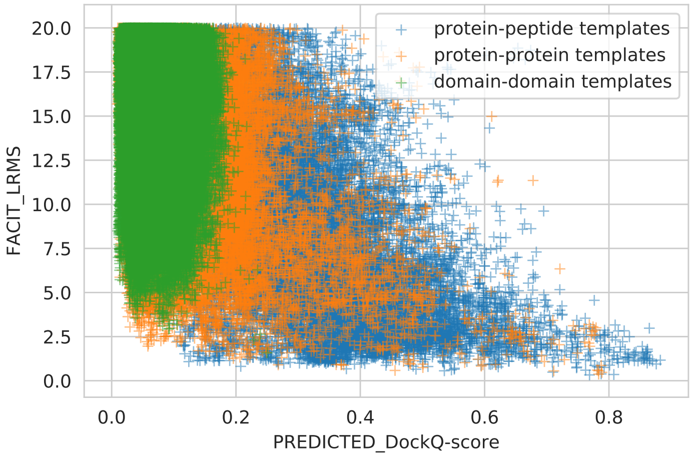


Figure S3: Distribution of LRMSD of different models versus their predicted DockQ-scores, as predicted by the random forest. Note the lack of sub-4 Å LRMS models derived from domain-domain templates. The figure has been cropped not to show results over 20.0 Å LRMSD. For ease of comparison and view, only 250,000 randomly selected models are shown for each kind of template.

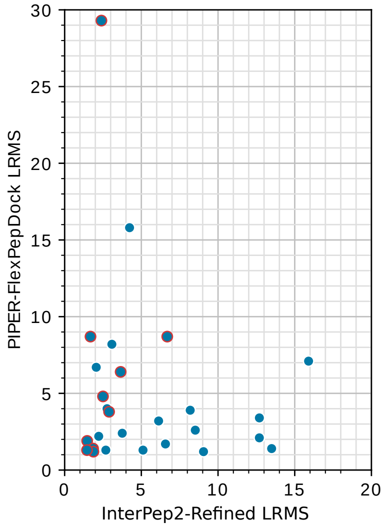


Figure S4: Comparison between results of InterPep-Refined and PIPER-FlexPepDock on the unbound set. The points with red edges denote which points would come from InterPep-Refined in the Combo method (InterPep2-PIPER-FlexPepDock). Thus, the LRMSD values for the Combo are on the x-axis for the points with a red edge and on the y-axis for the other points threshold on the InterPep2 score to decide whether to select that model or to select the PIPER-FlexPepDock model. In all but three cases the Combo methods selects the best possible model.
